# The absence of microbiome in germ-free mice exacerbates the pathogenicity of mouse-adapted SARS-CoV-2 MA10

**DOI:** 10.64898/2026.11.11.765867

**Authors:** EP Griffin, SH Feldman, R Jain, ST Le, MR Conaway, S Li, SR Robinson, J Larner, JW Mandell, JI Everitt, KL Andarawewa

## Abstract

Germ-free mice, which are devoid of microbiota, provide a powerful system to define the contribution of the microbiome to host responses during infection. Severe COVID-19 outcomes are associated with multiple comorbidities that are themselves characterized by microbiome dysbiosis; however, the causal role of the microbiota in SARS-CoV-2 pathogenesis remains poorly understood. Here, we use germ-free mouse models to evaluate the impact of the microbiome on disease severity following infection with mouse-adapted SARS-CoV-2 MA10. Comparison with conventionally housed mice revealed that the absence of the microbiome markedly exacerbates SARS-CoV-2–induced disease in both C57BL/6 and Swiss Webster backgrounds. Germ-free mice exhibited greater weight loss, impaired lung function, worsened pulmonary pathology, and pronounced alterations in circulating cytokine and chemokine profiles compared with conventional controls. These findings show that resident microbiota limit SARS-CoV-2 pathogenicity and establish germ-free COVID-19 models as a powerful platform to identify protective microbial taxa and microbiota-based interventions to reduce severe disease.

## Introduction

Germ-free (GF) mice, which are devoid of microbes, serve as a unique experimental system for studying the microbiome’s direct role in host responses during infection, as these models allow researchers to examine the consequences of microbial absence on disease progression. Dysbiosis of the microbiome has been associated with severe or fatal outcomes among COVID-19 patients (1). Yet the mechanisms by which commensal microorganisms influence COVID-19 severity remain incompletely understood. To address this significant gap in our knowledge, we characterized a GF mouse model of the mouse-adapted SARS-CoV-2-MA10 (MA10), which serves as a powerful tool for studying how the host microbiome influences COVID-19 pathogenesis.

## Results

To characterize the microbiome’s impact, we first utilized C57BL/6 (C57), one of the most commonly used mouse strains. GF and non-germ-free (NGF) mice were intranasally inoculated with MA10 virus (2). Mock-infected GF and NGF mice were compared with their respective infected groups. Subsequently, infected GF and NGF mice were also compared to determine the effect of microbiome absence on disease severity (Figure-1A).

**Figure 1.**
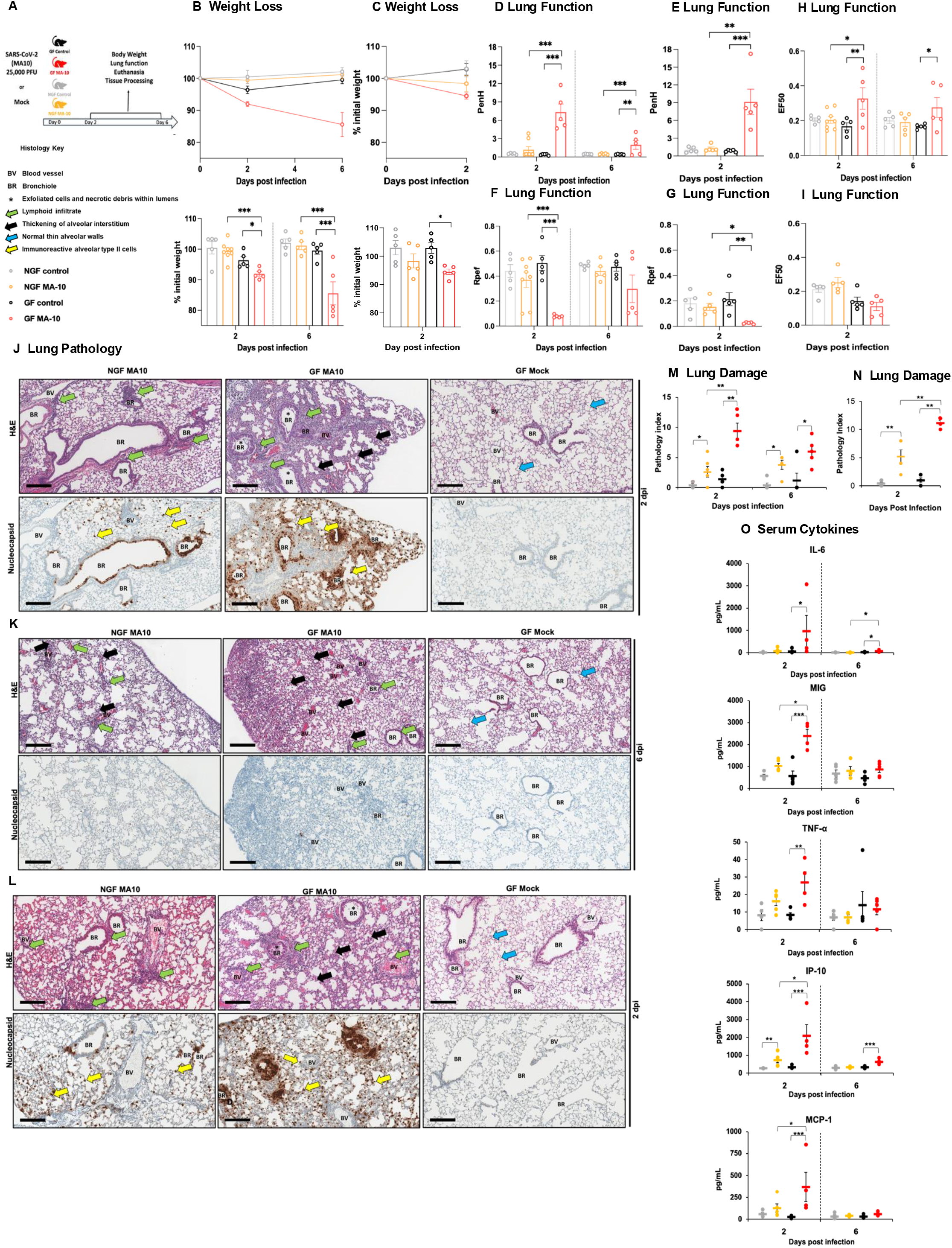
SARS-CoV-2 MA-10 infection is severely exacerbated in germ-free mice. (A) Experimental protocol. (B-C) Percent change in body weight: C57 (B) and SW (C). (D-I) WBP analysis of lung function parameters: C57 PenH (D), SW PenH (E), C57 Rpef (F), SW Rpef (G), C57 EF50 (H) and SW FE50 (I). (H-J) Representative images of lungs from mock and MA10-infected mice: C57 2 dpi (J), C57 at 6 dpi (K), and SW at 2 dpi (L). Top panels H&E and bottom SARS-CoV-2 nucleocapsid immunohistochemistry. (M-N) Lung damage: C57 (M) and SW (N). (O) C57 Serum cytokines. The box-and-whisker plot shows the mean±SEM. Each circle represents data for each mouse. *n*=4-5 mice per group. Scale bars: 200 μm. Where significant, *P* values are shown: ^*^*P*≤0.05; ^**^*P*≤0.01; ^***^*P*≤0.001. Swiss Webster:SW, and C57BL/6:C57.

All animals were monitored daily for clinical symptoms. Weight loss, lung function, lung and brain pathology, and serum cytokines were measured at 2 and 6 days post-infection (dpi).

Infected GF mice exhibited significantly greater weight loss than infected NGF mice at 2 and 6 dpi. Compared to their respective uninfected controls, both infected groups also showed significant weight loss (Figure-1B).

Lung function was measured by whole-body plethysmography (WBP) (3), including enhanced pause (PenH), relaxation time to peak expiratory flow (Rpef), and EF50, an indicator of expiratory flow rate to assess respiratory impairment. Infected GF mice exhibited a significant loss in pulmonary function than infected NGF mice, as indicated by altered PenH, Rpef, and EF50 values (Figure-1D, -F, -H).

The patterns of histologic changes in lungs were compared among the infected groups. At two dpi, many C57 NGF mice had minimal to mild bronchiolar epithelial cell degeneration and desquamation with debris in airway lumens (Figure-1J and S1A) and immune cell infiltrates in perivascular and peribronchial regions (Figure-S1B). Some NGF mice also had scattered areas of pulmonary parenchymal change characterized by intra-alveolar edema, accumulation of inflammatory cells, and hypercellular interstitial thickening of alveolar septal regions with immune cells. Alveolar parenchymal lesions were patchy in distribution and minimal to mild in severity in NGF mice (Figure-1J).

By 6 dpi airway epithelial changes were markedly reduced or absent (Figure-1K), with a slight increase in immune cell infiltration in peribronchiolar, perivascular, and alveolar regions. IHC analysis revealed that at 2 dpi, viral antigen was present in bronchial, bronchiolar, and alveolar type II epithelial cells (Figure-1J and S1C-D) and in the regions of virus-induced pulmonary injury. By 6 dpi, viral antigen was no longer present (Figure-1K).

In GF mice, at 2 dpi, MA10-induced pathology was similar in characteristics to that of NGF mice but more severe, with a greater viral antigen burden (Figure-1J and S2). By 6 dpi in GF mice, immunoreactivity for viral protein was lost, and airway epithelial pathology was markedly diminished (Figure-1K and S2). To quantify lung damage, a pathology index was generated by scoring bronchiolar epithelial changes, perivascular and peribronchial inflammatory infiltrates, and alveolar parenchymal inflammation and thickening (Figure-1M and supplemental material). The pathology index was significantly higher in the infected GF group than in the infected NGF group. No pathological changes or virus were detected in the brains of any animals (Figure-S3).

The severity of COVID-19 infection is linked to the presence of “cytokine storm.” Like in patients with moderate or severe COVID-19, compared with NGF mice, GF mice exhibited amplified inflammatory responses, with significantly elevated levels of pro-inflammatory cytokines and chemokines (Figure-1O), including IL-6, TNF-α, MCP-1, IP-10, and MIG (4).

Because host genetic background-dependent differences in disease susceptibility have been described for infectious diseases including influenza and SARS-CoV (5), we evaluated MA10 infection in Swiss Webster (SW) mice. Consistent with findings in C57 mice, GF SW mice inoculated at the same dose of virus exhibited increased weight loss (Figure-1C), greater impairment of lung function (Figure-1E, -G and -I), severe pulmonary pathology (Figure-1L and S2), increased viral antigen (Figure-1L), and a higher pathology index (Figure-1N and supplemental material) when compared to NGF SW mice.

## Discussion

Here, we modeled the pathogenesis of MA10 in GF mice and found that GF status exacerbates SARS-CoV-2 severity. When compared to NGF mice with a microbiome, GF mice exhibited significant changes in weight, lung function, lung pathology, as well as changes in cytokines/chemokines in the serum. Germ-free mouse models of SARS-CoV-2 may be useful as a model to identify beneficial gut microorganisms as a method to mitigate COVID-19 disease severity. Thus, GF mouse models are a robust platform for investigating microbiome–host interactions during SARS-CoV-2-MA10 infection.

It is important to note that in the pathology index we did not incorporate spontaneous pulmonary lesions (6) associated with the GF state and normal aging, such as pulmonary alveolar proteinosis (PAP) and lymphoid aggregates, as these findings were distinct from virus-induced pathology (Figure-S4). High viral immunoreactivity was noted in these areas of spontaneous lesions in GF mice. Identifying these baseline spontaneous lesions underscores the value of characterizing GF models not only for elucidating microbiome-dependent effects but also for revealing host-intrinsic features that may influence disease susceptibility or confound interpretation of study findings.

## Supplemental Figures

**Figure S1.**
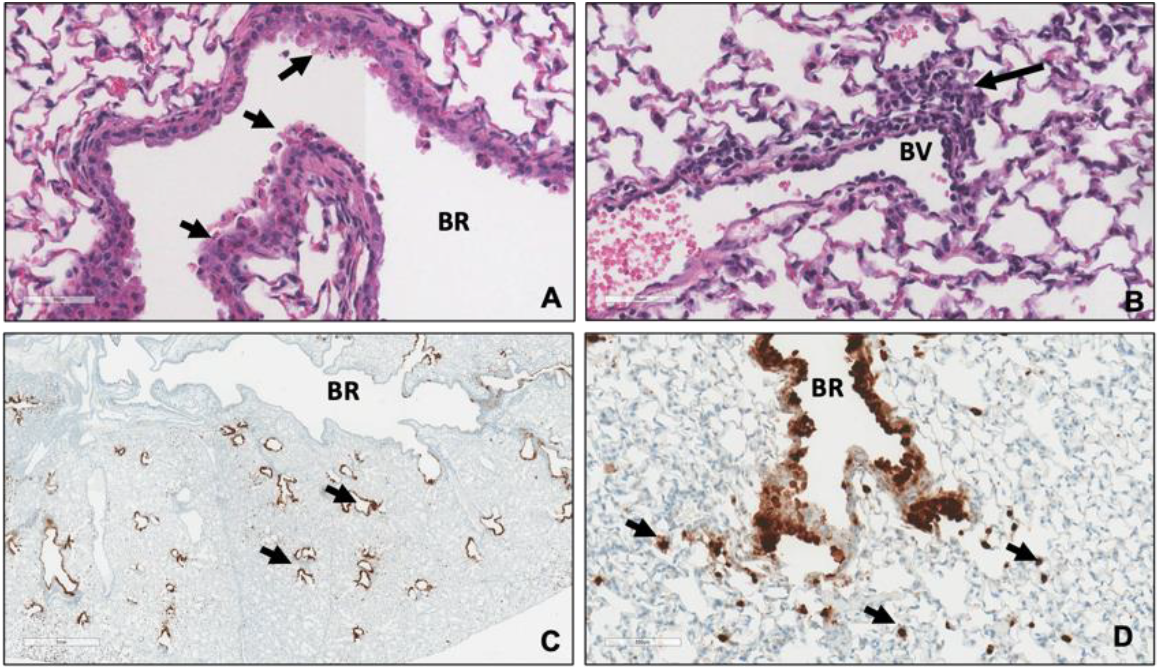
Photomicrographs of airways of MA10 infected NGF C57BL/6 mice at 2 dpi, related to Figure 1. (A) Changes in epithelium of terminal bronchiole (BR) with epithelial cell degeneration and sloughing (arrows) H&E Bar = 50μm. (B) Blood vessel (BV) with perivascular immune cell infiltrate (arrow) H&E Bar = 50μm. Note in both (A) and (B) lack of lesions in peribronchial and perivascular alveolar parenchyma. (C) Low magnification photomicrograph of lung lobe with strong immunoreactivity in all terminal bronchioles (arrows) and patchy strong immunoreactivity in mainstem bronchus (BR). SARS-CoV-2 immunostaining, Bar = 1mm (D) High magnification photomicrograph showing strong immunoreactivity in epithelial cells of terminal bronchiole (BR) as well as in surrounding alveolar type II epithelial cells (arrows) with normal adjacent pulmonary parenchyma. SARS-CoV-2 immunostaining, Bar = 100μm.

**Figure S2.**
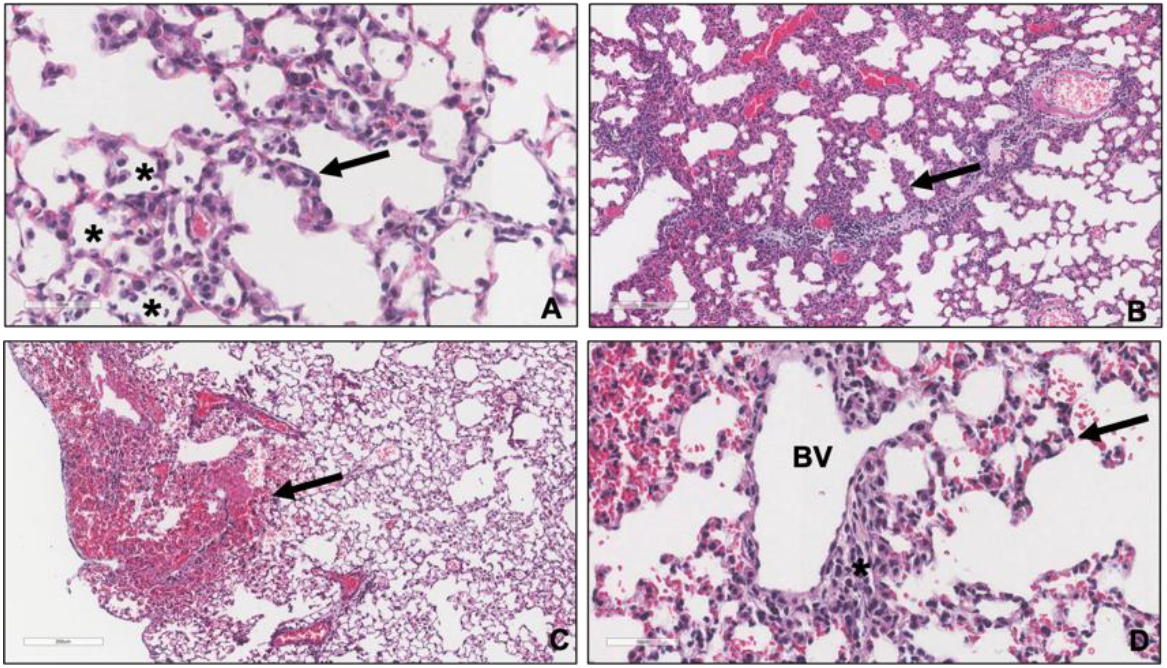
Photomicrographs of MA10-induced interstitial lung disease in GF mice, related to Figure 1. (A) Changes in alveolar region of infected C57BL/6 mouse 2 dpi showing cellular thickening of alveolar septal walls (arrow) and intra-alveolar accumulation of macrophages and inflammatory cells (^*^). H&E Bar = 50μm. (B) Lung changes in infected SW mice at 2 dpi showing interstitial cellular infiltration and thickening of alveolar parenchyma (arrow). H&E Bar = 200μm. (C) Low magnification photomicrograph of lung lobe with focus of hemorrhage (arrow) in a C57BL/6 mouse 6 dpi. H&E Bar = 200μm. (D) High magnification photomicrograph demonstrating interstitial infiltrate (arrow) in alveolar parenchyma extended from perivascular infiltrate (^*^) surrounding pulmonary blood vessels (BV) in a C57Bl/6 mouse 6 dpi. H&E Bar = 50μm.

**Figure S3.**
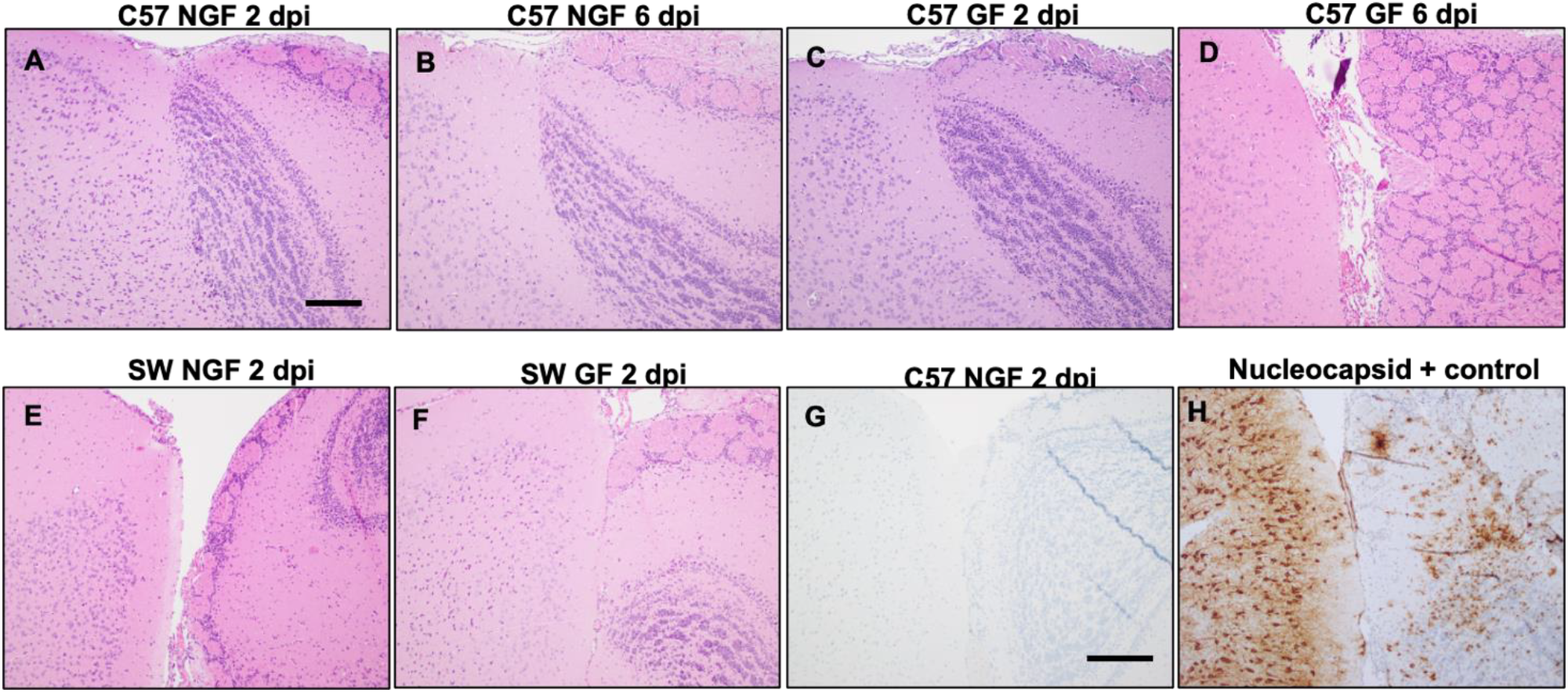
No evidence of histopathologic changes in brains of either GF or NGF mice, related Figure 1. All mouse brain images contain representative regions of interface between the cerebral cortex on the left half (prelimbic) and the olfactory bulb on the right. (A-F) No morphologic evidence of neuronal ischemic injury, necrosis, nor apoptosis was detected. No gliosis, acute or chronic inflammatory infiltrates were identified in brain parenchyma or meninges. Scale bar in A represents 100 microns, and applies to A-F. Lack of SARS-CoV-2 nucleocapsid protein immunoreactivity in brains of GF or NGF mice, related to Figure-1. (G) A NGF mouse at 2 dpi reveals no SARS-Co-V-2 nucleocapsid protein immunoreactivity. This was observed in all mouse models described in the manuscript. (H) As a positive control, a humanized (AC70 transgenic) mouse brain at 4 dpi shows abundant SARS-CoV-2 Nucleocapsid protein immunoreactivity in both cerebral cortex (left half) and olfactory bulb (right half). Scale bar in G represents 200μm and applies to G and H.

**Figure S4.**
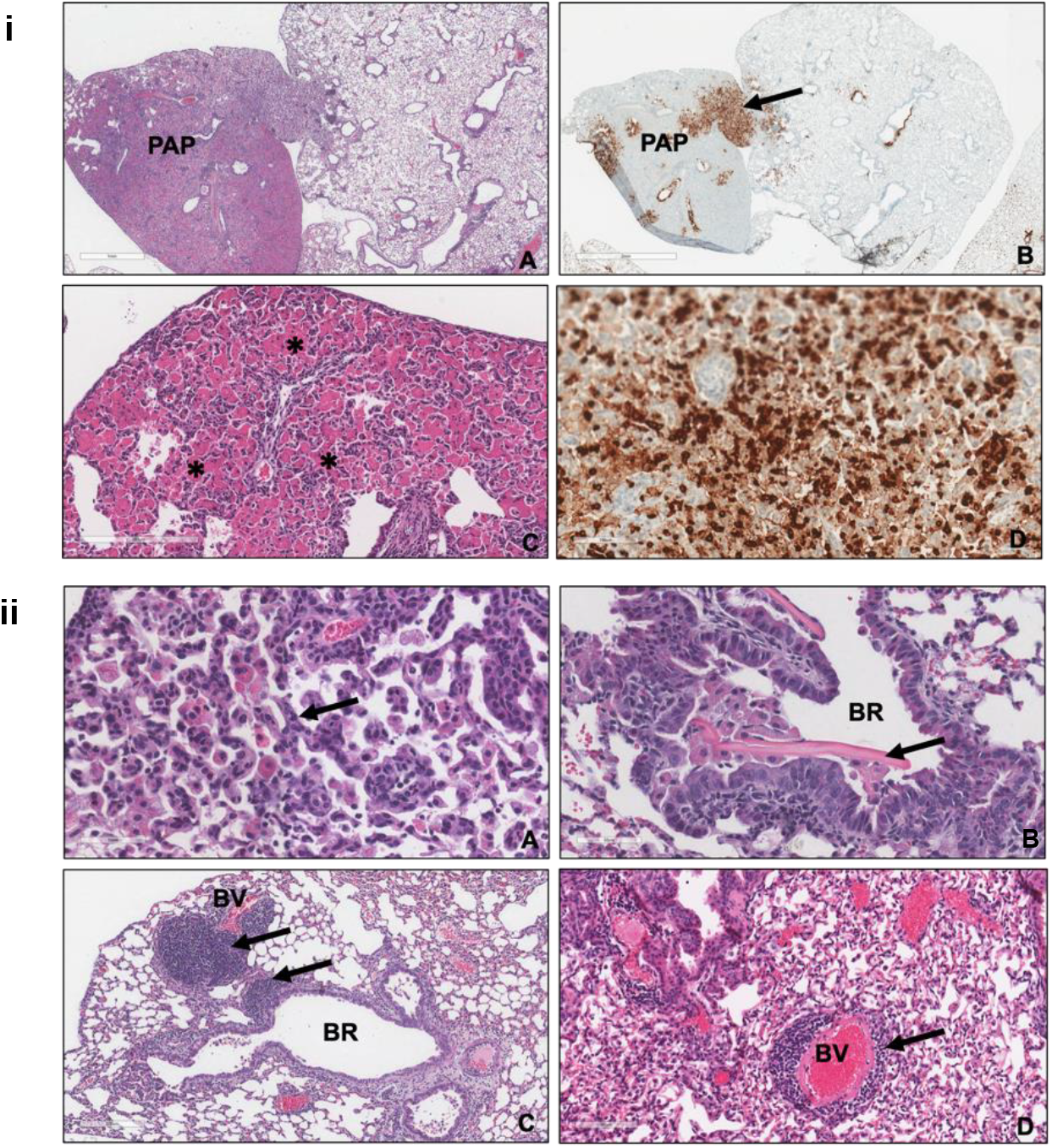
S4i. Photomicrographs of PAP in GF C57BL/6 mice infected with MA10, related to Figure 1. (A) Low magnification photomicrographs show lobar consolidation with PAP with admixed lymphoid infiltrate. H&E Bar = 1mm. (B) Immunostaining for SARS-COV-2 protein in PAP region identical to (A) depicting strong patchy staining (arrow) 2 dpi. Bar = 2mm. (C) Photomicrograph of showing intra-alveolar accumulation of eosinophilic proteinaceous material and macrophages (^*^) characteristic of PAP. H&E Bar = 200μm. (D) High magnification photomicrograph of immunostaining in (B) showing intense immunoreactivity in alveolar type II epithelial cells 2 dpi. Bar = 100μm. **S4ii. Examples of spontaneous pulmonary pathology associated with age and GF status in study mice, related to Figure 1**. (A) High magnification photomicrograph of region of PAP in uninfected GF SW mice. Note hypertrophic and hyperplastic alveolar type II epithelial cells (arrow) lining alveoli filled with large mono- and binucleate macrophages filled with abundant eosinophilic cytoplasm. H&E Bar = 50μm. (B) Terminal bronchiole in GF uninfected C57BL/6 mouse containing large crystalloid eosinophilic structure in the region of PAP. H&E Bar = 50μm. (C) Lung photomicrograph in C57BL/6 GF mouse 2 dpi MA-10 demonstrating large lymphoid nodules in peribronchial (BR) and perivascular (BV) regions considered spontaneous background changes. H&E Bar = 200μm. (D) Photomicrographs of untreated control SW mice demonstrating prominent perivascular (BV) lymphoid cuffing (arrows). H&E Bar = 100μm.

## Supplemental material

## Materials and Methods

### Sex as a biological variable

Data were obtained from both male (n = 3) and female (n = 60) animals. Sex was recorded at the time of data collection but not analyzed separately, as sex-based differences were not the study’s main focus.

### Mouse experiments

Nine to eighteen month-old specific pathogen free (SPF) Swiss Webster and C57BL6 mice (Taconic Labs), and GF Swiss Webster and GF C57BL6 mice (University of Virginia Germ Free Core Facility) were used for these experiments. The GF mice were bred and maintained in flexible film, positively pressured isolators (Class Biological Clean) at the University of Virginia, Gnotobiotic Mouse Facility. The GF mice were transported in specialized autoclaved biocontainment cages and were housed at the University of Virginia Biosafety Level 3 (BSL3) facility. Anesthetized (Isoflurane or Ketamine/Dexmedetomidine 50-150 mg/kg/0.25-0.5mg/kg or Ketamine/Xylazine 60-80mg/kg/5-10mg/kg) mice were inoculated intranasally with 25,000 PFU SARS-CoV2 MA10, and 50ul of Vero-E6 DMEM/10% FBS medium was used as a mock-infection for control animals. The infected mice were observed once daily to monitor clinical symptoms. Body weights were obtained prior to start of the experiment and at the end of the experiment on 2 or 6 dpi. Lung function was assessed utilizing whole body plethysmography (WBP; DSI Buxco respiratory solutions, DSI Inc.) at indicated time points by allowing mice to acclimate in WBP chambers for 30 minutes, followed by 5 minutes of data recording as described previously (3). Acquired data was analyzed using FinePointe software. Mice were euthanized at indicated time points or analysis. All procedures involving infectious virus were approved by the University of Virginia Institutional Biosafety Committee, and were conducted in ABSL3 facility using approved practices. All animal procedures were approved by the University of Virginia Animal Care and Use Committee.

### Viruses and cells

SARS-CoV-2 MA10 was obtained from BEI resources. All virus stocks were propagated on Vero E6 (ATCC) cells in Dulbecco′s Modified Eagle′s Medium (Gibco) and supplemented with 10% fetal bovine serum (Gibco), 1% Antibiotic-Antimycotic mix (Gibco), and incubated at 37 °C, 5% CO2. Viral titers of the supernatant were determined using a standard plaque forming assay. All viral propagations and infections were conducted at BSL-3 facility.

### Cytokine and chemokines analysis

Mouse cytokine 32-plex assay (Millipore Sigma Cat. No. MCYTOMAG-70K) was utilized to analyze chemokines and cytokines in serum from mice according to the manufacturer’s protocol (with a 12h incubation period). Cytokines were recorded on a MAGPIX machine (Luminex) and the analysis was done on MILLIPLEX Analyst software Version 5.1.0.0.

### Immunohistochemistry and H&E Staining

Tissues were fixed in 10% neutral buffered formalin and processed by the UVA Research Histology Core on a Shandon Excelsior processor, embedded in paraffin using a Histocore embedding station. Blocks were sectioned at 5 microns on a Leica RM2255 microtome and placed on Fisher Brand superfrost plus slides. H&E was done on an automated stainer, Gemini Varistain. Immunohistochemistry staining for SARS-CoV-2 nucleocapsid was performed by the University of Virginia Biorepository and Tissue Research Facility on a robotic platform (Ventana Discover Ultra Staining Module, Ventana Co., Tucson, AZ, USA). A heat-induced antigen retrieval protocol set for 64 min was carried out using a TRIS–ethylenediamine tetraacetic acid (EDTA)–boric acid pH 8.4 buffer (Cell Conditioner 1). Endogenous peroxidases were blocked with peroxidase inhibitor (CM1) for 8 min before incubating the cells with SARS-CoV2 nucleocapsid antibody (ProSci Cat# 9099) at 1:2,000 dilution for 60 min at room temperature. Antigen-antibody complex was then detected using DISC. OmniMap anti-rabbit multimer RUO detection system and DISCOVERY ChromoMap DAB Kit (Ventana Co.). Subsequently, all the slides were counterstained with hematoxylin; they were dehydrated, cleared, and mounted for the assessment.

### Pathology index

The lung pathology of MA10-infected mice was assessed and scored by a board-certified veterinary pathologist (JIE) with knowledge of the allocation group initially to determine the extent and type of lesions present due to the presence of spontaneous pulmonary pathology associated with the GF state. Viral pathology of MA10-infected and control lungs were scored in a masked fashion without the observer’s knowledge of the allocation group for three separate parameters: bronchiolar epithelial changes, perivascular and peribronchial inflammatory infiltrates, and alveolar parenchymal inflammation and thickening. A pathology index was generated for each mouse by totaling the three scored parameters. Spontaneous pathology associated with the GF state and normal aging (such as increased bronchus-associated lymphoid tissue: (BALT), perivascular and peribronchial lymphoid cuffing, and PAP) were not included in the pathology index scoring of viral-induced changes. Due to the lack of neutrophils as a dominant part of the inflammatory infiltrate, and due to the relative paucity of fibrin and diffuse alveolar damage noted, the study pathologist decided not to utilize the acute lung injury (ALI) index and diffuse alveolar damage (DAD) index utilized in some previous MA10 studies (2).

### Statistical analysis

Two-way analysis of variance was used for the analysis of the weight and plethysmography outcomes. Contrasts were used for specific comparisons among groups.

Pathology scores were compared with the nonparametric Mann-Whitney test. Two-way ANOVA was also used for the analysis of the cytokine levels after transforming to the log-scale, in order to stabilize the variance as well as to facilitate interpretation as fold changes across groups. The cytokine p-values displayed are based on comparisons after transforming the cytokine levels to the natural log scale. Analyses were carried out in SAS version 9.4 and GAUSS version 22.

### Study approval

Animal experiments were approved by the IACUC of the University of Virginia. SARS-CoV-2 experiments were approved by the University of Virginia Institutional Biosafety Committee.

## Data availability

All data in this study are available and are shown as individual data points on the figures.

## Acknowledgments

This work was supported by a grant to KLA from The Ivy Foundation, COVID-19 Translational Research Fund (#IVY13).

The authors would like to thank the Center for Comparative Medicine for its support, especially Rory Laman, Susan Landes, and Gina Wimer, for their assistance with animal work.

The authors also acknowledge the support of the Biorepository and Tissue Research Facility, which is supported by the University of Virginia School of Medicine, Research Resource Identifiers (RRID): SCR_022971.

The cytokine data for this manuscript were generated in the University of Virginia Flow Cytometry Core Facility (RRid:SCR_017829) and is partially supported by the NCI Grant (P30-CA044579).

The authors acknowledge the New England Regional Biosafety Laboratory at Tufts University and support from NIH UC7 AI180310 “Resources and workforce development for the New England Regional Biosafety Laboratory”.

## Author Contributions Statement

EPG performed experiments, acquired, analyzed, and interpreted data, generated figures, and drafted the manuscript; SF conducted experiments, analyzed data and revised manuscript; RJ and STL acquired and analyzed data; MC analyzed the data and drafted the manuscript; SL contributed to generating figures and edited the manuscript; SRR contributed to generating figures; JL guided the individual experiments; JWM analyzed and generated figures; JE analyzed data, generated figures, and drafted and revised manuscript; KLA conceived, designed, and performed experiments; supervised the project, secured funding, analyzed and interpreted the data; drafted and revised the manuscript.

